# Climate-Driven Ecosystem Productivity Changes Restructure Food Systems

**DOI:** 10.1101/2025.10.14.682485

**Authors:** Hamid Dashti, Meng Luo, Charles Nicholson, Min Chen

## Abstract

Climate change affects food systems through multiple pathways, yet the isolated effects of climate-driven ecosystem productivity changes remain poorly understood. While field studies translate productivity changes to crop yields, at larger scales yield conflates productivity with management and technology, and critically, productivity changes occur across all vegetation types, not just agricultural lands. We isolate ecosystem productivity changes to trace their propagation through food systems across three socioeconomic scenarios (SSP1-2.6, SSP3-7.0, SSP5-8.5) by coupling the CLASSIC land surface model with GCAM integrated assessment model. Productivity changes lead to crops systematically shifting from human consumption to animal feed and bioenergy (up to 15% reallocation), while there is a significant transition of managed agricultural lands to unmanaged lands by up to 6 million km^2^ as productivity gains enable less intensive land use. These changes drive extreme regional divergence, with food production volume changing from +35% to -28% and substantial portfolio reorganization. Disruptions peak under regional rivalry (SSP3-7.0), revealing how fragmented governance amplifies biophysical feedbacks beyond climate forcing alone. While how much we eat may remain stable, ecosystem productivity fundamentally reshapes what we eat and where to source it.

## 1 Introduction

Climate change is fundamentally reshaping global food systems through multiple destructive pathways, from temperature stress to pathogens and extreme weather events (FAO 2018; Wheeler and von Braun 2013; Hultgren et al. 2025; Yang et al. 2024; Ristaino et al. 2021). Yet widespread observations and modeling studies show that climate change simultaneously drives vegetation productivity surges in many regions through factors like CO2 fertilization and warming (Li et al. 2024; Piao et al. 2020). This productivity surge appears unambiguously beneficial for agriculture and food production, but how these ecosystem changes cascade through markets, land use, and regional systems remains a critical blind spot in food system projections.

From a physiological perspective, climate change affects ecosystem productivity through net primary production (NPP), the carbon captured via photosynthesis (Piao et al. 2020). In field studies of food systems, these NPP changes are typically translated to crop yields (Hultgren et al. 2025; Zhang et al. 2020; Srinivasan et al. 2017), providing crucial insights into physiological mechanisms. However, when projecting food systems at larger scales, yield and productivity become separate and potentially divergent outcomes. First, yield inherently bundles ecosystem productivity responses with management practices, technological advancement, and adaptation strategies (Potapov et al. 2022; Lobell et al. 2002), making it difficult to separate Earth system feedbacks from socioeconomic interventions. Second, climate-driven NPP changes occur across all terrestrial ecosystems including unmanaged lands like natural forests and grasslands, not just agricultural systems. Previous studies show that forest productivity changes significantly alter land use patterns(Luo et al. 2025), but how these effects ripple through food systems remains unknown.

These productivity-driven changes play out differently depending on which development pathway society takes. The shared socioeconomic pathways coupled with representative concentration pathways (SSP-RCPs) provide the standard framework for exploring these futures (Riahi et al. 2017). While the Paris Agreement envisions sustainable development to limit warming below 2°C (SSP1), recent geopolitical disruptions such as pandemics, trade wars, and regional conflicts suggest we are moving toward futures characterized by regional rivalry (SSP3) or fossil-fueled growth (SSP5) (Patel and Hansmeyer 2024). Each pathway incorporates distinct strategies for agricultural transformation, adaptation, and land use governance that fundamentally shape how productivity gains propagate through food systems.

Here we use the Global Change Analysis Model (GCAM) integrated assessment model (Calvin et al. 2019) coupled with the CLASSIC land surface model (Swart et al. 2019) to dynamically update vegetation productivity in GCAM, which assumes fixed productivity. We conducted counterfactual experiments comparing food system trajectories with and without dynamic vegetation feedbacks across three socioeconomic pathways (SSP1-2.6, SSP3-7.0, and SSP5-8.5). This design isolates the marginal effects of climate-driven productivity changes from baseline technological advancement and management, revealing how biophysical feedbacks alone propagate through global food systems.

We find that higher productivity doesn’t create food abundance but instead triggers fundamental reorganization of both food quantity and composition: crops systematically flow from direct human consumption to animal feed and bioenergy uses (up to 15%), while food systems produce more with dramatically less managed land. This reshuffling creates extreme regional divergence with food system capacity changes ranging from +35% to -28%. Disruptions are smallest under sustainable development (SSP1-2.6) but peak under regional rivalry (SSP3-7.0).

## 2 Method

We employ a counterfactual modeling approach to isolate the marginal effects of climate-driven vegetation productivity changes on global food systems. By comparing GCAM simulations with and without dynamic vegetation productivity updates, we quantify how biophysical feedbacks alone, separate from baseline technological and management trends, propagate through agricultural markets, land use decisions, and food allocation. The vegetation productivity updates are derived from the CLASSIC land surface model forced by Earth System Model climate projections under three socioeconomic pathways (SSP1-2.6, SSP3-7.0, SSP5-8.5), representing a soft one-way coupling.

### 2.1 GCAM Baseline Configuration

GCAM is a widely used integrated assessment model that represents interactions among economy, energy, water, land, and climate systems across 32 geopolitical regions (Calvin et al. 2019). For food systems, GCAM simulates market equilibrium where agents make resource allocation decisions based on prices, costs, supply, and demand across agricultural and livestock sectors (EDMONDS et al. 2017). The model represents 15 crop types and 5 livestock products. GCAM’s climate component, Hector, is a reduced-complexity model that calculates radiative forcing and temperature from greenhouse gas emissions but maintains vegetation and soil carbon densities at fixed historical levels without dynamic climate feedbacks (Dorheim et al. 2023). This static carbon assumption, where carbon densities remain constant regardless of climate change, represents the baseline condition we modify through our coupling with CLASSIC.

### 2.2 Vegetation Productivity Coupling Framework

We implement dynamic vegetation feedbacks in GCAM using productivity outputs from the CLASSIC land surface model (Swart et al. 2019) obtained through the Inter-Sectoral Impact Model Intercomparison Project Phase 3b database (ISIMIP3b, 2024). CLASSIC was forced with meteorological data from the Geophysical Fluid Dynamics Laboratory Earth System Model version 4 (GFDL-ESM4, Dunne et al. 2020), providing net primary productivity (NPP) and heterotrophic respiration (HR) projections under three climate scenarios. We selected NPP and HR as proxy variables because they isolate the productivity response to climate forcing without the confounding effects of disturbance history and legacy carbon that affect stock measurements (Luo et al. 2025). While these metrics do not account for all factors affecting vegetation change such as insect damage or forest management shifts, they provide consistent indicators of climate-driven productivity responses across regions and time periods. The full coupling methodology is detailed in Luo et al. 2025.

The coupling translates CLASSIC outputs into carbon density updates for GCAM through scalar calculations (Bond-Lamberty et al. 2014; Thornton et al. 2017). Vegetation carbon scalars are calculated as the ratio of projected NPP to historical mean NPP (5-year average in 2010–2014) for each region. Soil carbon scalars incorporate both NPP changes, which increase soil carbon through litter inputs, and HR changes, which decrease soil carbon through decomposition. These gridded annual scalars are aggregated to GCAM’s 384 subregions using area-weighted averaging at 5-year intervals. After harmonizing plant functional type classifications between CLASSIC and GCAM, we retrieve baseline carbon densities, apply the calculated scalars, and update GCAM’s land carbon density (C/m^2^) every five years. We also apply vegetation carbon scalars to the yields of managed crops, bioenergy, managed pasture, and managed forests, capturing how productivity changes affect agricultural output beyond just carbon storage (Luo et al. 2024).

### 2.3 Counterfactual and Scenario Design

We conduct counterfactual analyses comparing GCAM simulations with and without vegetation productivity coupling across three socioeconomic development pathways: SSP1-2.6, SSP3-7.0, and SSP5-8.5. For each SSP-RCP combination, we run two simulations: a baseline with static carbon densities reflecting standard GCAM configuration, and a treatment incorporating dynamic vegetation updates from CLASSIC.

We examine three contrasting futures that span the range of plausible development trajectories. SSP1-2.6 represents sustainable development with strong international cooperation and aggressive climate mitigation to limit warming below 2°C, featuring rapid technology transfer, dietary shifts, and forest protection. SSP3-7.0 depicts regional rivalry with nationalistic policies, slow economic development, and material-intensive consumption, where countries prioritize domestic food security over global cooperation. SSP5-8.5 assumes fossil-fueled development with rapid economic growth, high technological progress, and intensive resource exploitation, representing a world that prioritizes development over environmental protection (Riahi et al. 2017; Patel and Hansmeyer 2024). These pathways embed fundamentally different assumptions about agricultural technology, land use policies, trade openness, and bioenergy deployment that shape how vegetation productivity changes propagate through food systems.

### 2.4 Food System Change Metrics

We quantify vegetation-driven food system changes through three complementary metrics that capture production, composition, and allocation responses. Food system absolute change measures the total production response as ΔProduction = Production_updated_ − Production_baseline_ in crop and livestock production (Mt) for each region and scenario. We also calculate percentage change relative to baseline. Portfolio reorganization quantifies compositional shifts within food systems as 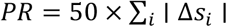 where 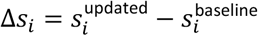 represents the change in production share of food item *i*. This metric scales from 0 (no change in composition) to 100 (complete reorganization). Portfolio reorganization is calculated separately for crops and livestock, then the two category scores are averaged to obtain total portfolio reorganization, revealing whether regions maintain traditional agricultural patterns or shift to entirely different production mixes under vegetation forcing.

To assess demand-side responses, we track percentage point changes in crop allocation across six end-use categories: food staples, food non-staples, animal feed, ethanol, biomass oil, and industrial uses. For each crop type, we calculate the share of total demand flowing to each end-use in baseline versus coupled simulations, with the difference representing reallocation induced by vegetation productivity changes. Positive values indicate vegetation productivity updates lead to crops shifting toward that use, while negative values show crops moving away from it. All metrics are calculated at 5-year intervals from 2015 to 2100 across GCAM’s 32 regions, enabling assessment of both temporal evolution and regional heterogeneity in food system responses.

## 3 Results

Vegetation productivity changes fundamentally reorganize global food systems rather than simply boost production. While total food production increases from 2015 to 2100 under all scenarios to meet growing population demand, incorporating dynamic vegetation feedbacks creates substantial marginal changes beyond baseline technological and demographic trends. These vegetation-driven impacts transform what is produced, where it is produced, and how it is allocated across competing uses, with development pathways determining the scale of disruption.

### 3.1 Vegetation productivity drives divergent commodity responses regionally and globally

At the global level, development pathways shape the magnitude of reorganization (Figure1). Sustainable development (SSP1-2.6) maintains relative stability with modest changes across food types, while regional rivalry (SSP3-7.0) and fossil-fueled growth (SSP5-8.5) trigger substantial redistributions. Figure 1 reveals sharp commodity-specific contrasts: soybean (+188 Mt) and oil crops (+146 Mt) surge while sugar crops (-183 Mt) and corn (-174 Mt) collapse. Livestock sectors respond differently, with all animal products remaining stable or increasing slightly (<2%), and dairy showing the largest absolute gains. This divergence between volatile crop responses and stable livestock production suggests that vegetation feedbacks affect plant and animal systems through fundamentally different mechanisms.

**Figure 1.**
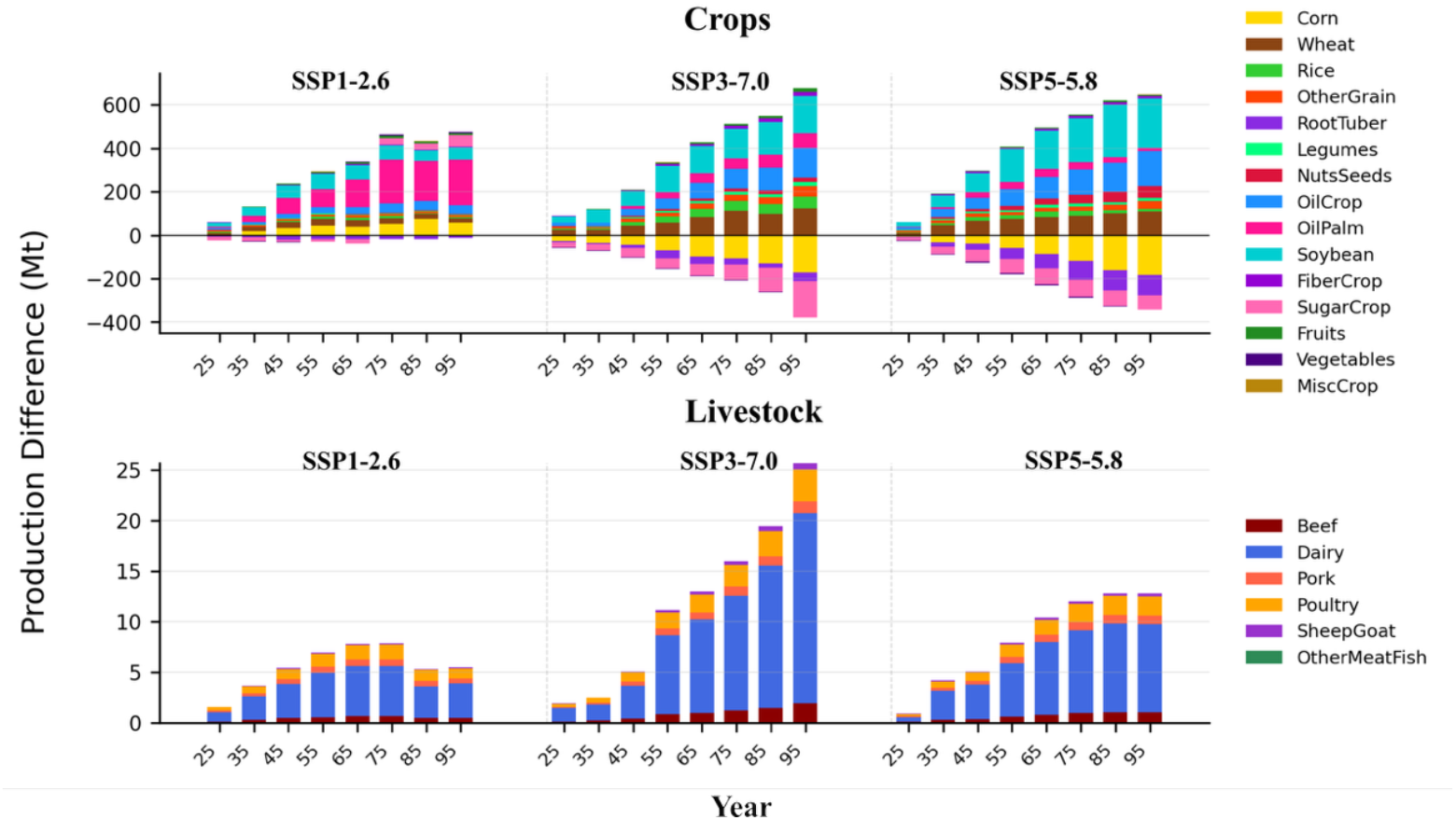
Global production changes (Mt) by commodity under vegetation-climate coupling across three SSP scenarios, 2100.

Vegetation productivity changes trigger highly heterogeneous regional responses, with individual regions experiencing production changes from +35% to -28% and major shifts in crop portfolios. Figure 2 displays the 15 regions with the most extreme changes in individual commodity production across all SSPs and food types. Brazil experiences the largest absolute losses across all SSPs, declining by 74 to 230 Mt depending on the scenario. Australia-New Zealand follows a similar pattern with losses of 23 to 93 Mt. In contrast, EU-15 gains between 47 and 152 Mt while Canada shows comparable increases. Generally, SSP3-7.0 and SSP5-8.5 lead to the largest disruptions, though exceptions occur. Indonesia, for example, gains over 100 Mt under SSP1-2.6 (almost entirely from oil palm) compared to just 10 Mt of total food production under SSP5-8.5.

**Figure 2.**
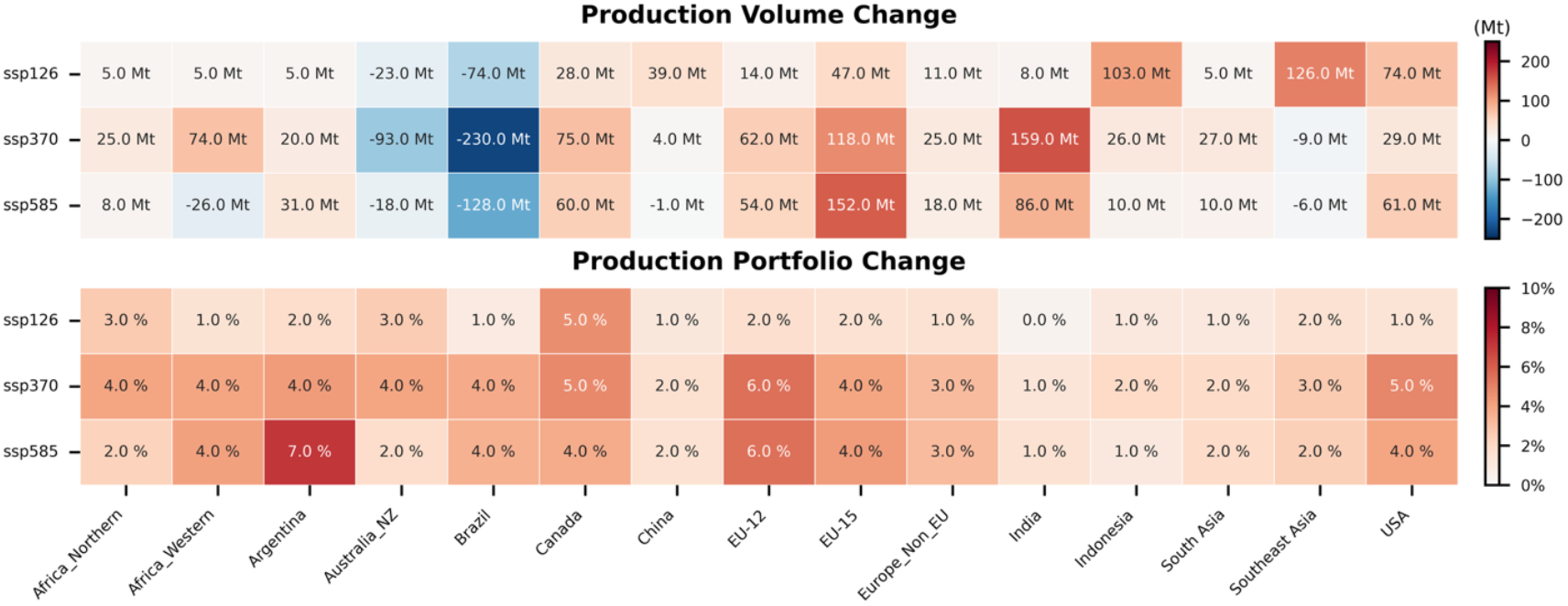
Production volume and portfolio changes in most-affected regions. Regions selected for having the largest change in any individual commodity under vegetation coupling. Shows both total production changes (Mt) and agricultural portfolio reorganization across three SSP scenarios in 2100.

These volume changes coincide with varying degrees of portfolio reorganization (Figure 2). Brazil’s losses concentrate heavily in sugar crops, which account for 85% of crop losses under SSP3-7.0, while the Australia-New Zealand region shows losses distributed more evenly across wheat, sugar crops, and dairy (Figure S2). This pattern of concentrated versus distributed impacts appears throughout regions, revealing how vegetation productivity feedbacks can drive single-commodity surges (Indonesia’s oil palm), single-commodity collapses (Brazil’s sugar), or broad-based restructuring across multiple agricultural products.

### 3.2 Managed-to-Unmanaged Land Transitions

Dynamic vegetation productivity drives systematic conversion from managed to unmanaged land systems globally (Figure 3). Managed lands retreat by 891 thousand km^2^ under SSP1-2.6 but exceed 6,000 thousand km^2^ losses under SSP3-7.0 and SSP5-8.5—a seven-fold increase. Unmanaged systems expand correspondingly. Cropland shows the largest absolute declines, losing up to 2,592 thousand km^2^ (20% reduction), followed by managed hardwood forests (1,160 thousand km^2^). Conversely, unmanaged pastures expand by 2,348 to 2,560 thousand km^2^ under higher emission scenarios.

**Figure 3.**
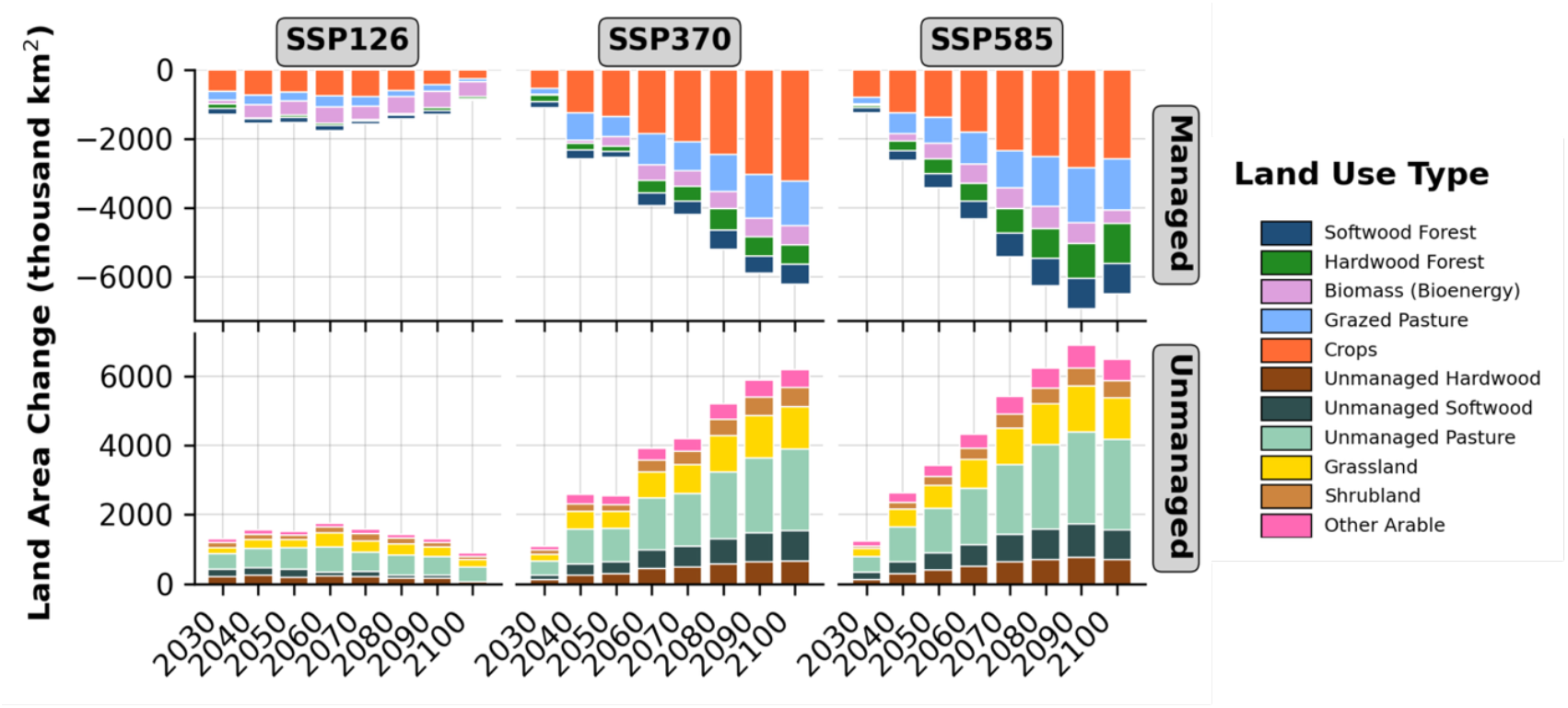
Global land transitions (thousand km^2^) from managed to unmanaged systems under vegetation-climate coupling across SSPs, 2100.

Similar to food, regional land transitions show substantial heterogeneity as well (Figure S5). Russia, China, and USA consistently experience the largest changes across all scenarios. In some regions, land use changes align directly with food production outcomes. Brazil loses 107 thousand km^2^ of cropland and 186 thousand km^2^ of grazed pasture, corresponding with its 74-230 Mt food production decline. However, other regions show more complex patterns. The USA under SSP5-8.5 loses 212 thousand km^2^ of cropland and 413 thousand km^2^ of managed forests, yet experiences a net food gain, producing 100 Mt more soybean despite losing 70 Mt of corn. These contrasting patterns indicate that vegetation feedbacks create diverse land-food relationships across regions, from straightforward declines to paradoxical reorganizations where production increases despite land losses.

### 3.3 Crops systematically shift from food to fuel and feed uses

Vegetation productivity increases trigger systematic reallocation of crops away from human consumption (Figure 4). Oil crops experience the most dramatic shifts, losing 9.4 percentage points from food uses while gaining 15.0 percentage points for bioenergy under SSP3-7.0. SSP5-8.5 shows similar patterns with 5.6 percentage point losses from food and 10.8 percentage point gains for bioenergy, while SSP1-2.6 exhibits smaller changes. Wheat and rice consistently shift from staple foods to animal feed across all scenarios; wheat loses 5.4 percentage points from food while gaining 6.9 percentage points for feed under SSP3-7.0, and rice shows similar patterns. Soybean reallocations split between bioenergy gains (3.8-4.7 percentage points) and losses from both food and feed, while corn shows variable responses across scenarios. Generally, SSP3-7.0 exhibits more significant reallocations compared to other pathways. These demand reallocations complete the reorganization picture: vegetation feedbacks simultaneously alter what is produced (Figures 1-2), how land is used (Figure 3), and where crops ultimately flow (Figure 4), with changes in each domain reinforcing the others.

**Figure 4.**
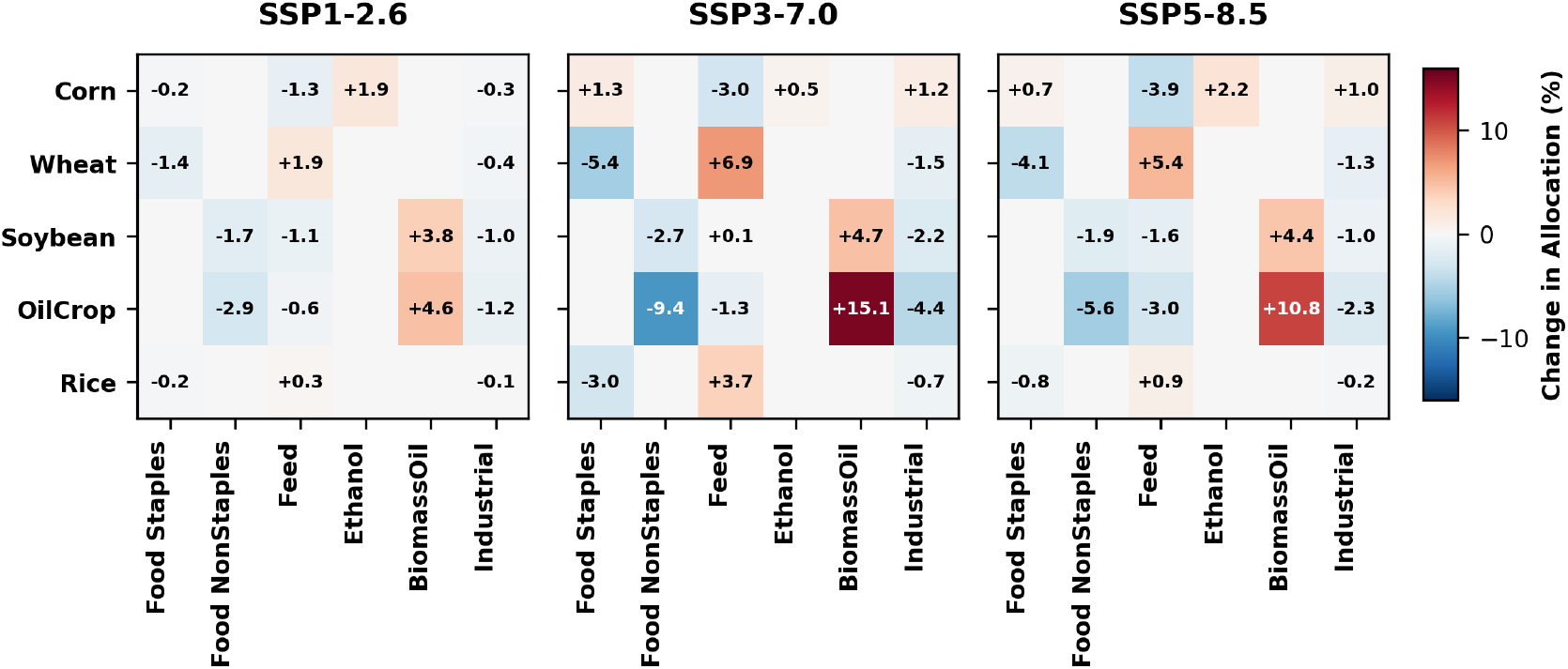
Percentage point changes in crop allocation across end-uses (food, feed, bioenergy, industrial) under vegetation-climate coupling, 2100.

## 4 Discussion

Decades-long field-scale experiments such as FACE studies have long established that climate factors such as CO2 fertilization enhance productivity of C3 crops and under certain conditions C4 crops, but this productivity is dampened by other climate factors such as heat stress or drought (Ainsworth and Long 2021; Lesk et al. 2016). Our counterfactual approach isolates climate-driven vegetation productivity changes from these confounding physiological factors, as well as from management practices and technological advancement that play significant roles in global food systems. Critically, our approach captures productivity changes across all terrestrial vegetation, not just agricultural lands. We show that rather than food simply becoming more abundant, productivity improvements lead to systematic food system reorganization driven by regional and sectoral competition, land use change, and market dynamics that redirect crops from human consumption toward bioenergy and feed uses. While climate change in reality involves multiple simultaneous impacts, isolating individual pathways provides deeper understanding of physiological-socioeconomic interactions.

The central mechanism driving food system reorganization operates through differential capacity for demand expansion across agricultural end-uses. When vegetation productivity increases generate surplus agricultural output, human food consumption cannot absorb this surplus because caloric requirements remain relatively stable (∼2000-4000 calories per capita; Sands et al. 2023). This demand saturation redirects surplus production toward sectors with greater capacity for expansion, particularly animal feed and biomass oil production for biofuels (Figure 4). The systematic crop substitution follows this expansion capacity, where corn allocation to bioenergy remains minimal while soybean and oil crop allocation to biomass oil increases dramatically. This reallocation pattern drives substantial production shifts, with biomass oil feedstocks (soybeans and oil crops) increasing by 794 Mt total across scenarios while ethanol feedstock (corn) decreases by 327 Mt. Land allocation reorganizes accordingly, with managed agricultural lands retreating as productivity gains allow less intensive land use to meet market demands, maintaining the supply-demand balance central to GCAM’s economic optimization. However, these global patterns manifest with severe regional differences, ranging from +35% to -28% changes in food system capacity, with regional rivalry (SSP3-7.0) amplifying disruptions most severely, reflecting how fragmented governance intensifies competition between food and non-food agricultural uses.

The implications of these agricultural reorganizations change what we eat and where we get it rather than how much we consume overall. While global food production increases, the extreme regional variations in food systems, mean that local food availability and composition may shift dramatically in many regions. Under SSP370, regions experiencing productivity-driven declines in food capacity cannot easily access surplus production from more productive regions due to fragmented governance and trade barriers. This may create a two-tiered system where wealthier regions can afford to import needed food products while poorer regions are forced to adapt their diets based on whatever their domestic production systems can provide under the new agricultural reorganization. The systematic withdrawal of crops from direct food uses toward biofuel production compounds these regional disparities, potentially making certain food categories less available in regions that lose comparative advantage. This is in line with other studies that show SSP3 leads to significant disruption of food systems (van Meijl et al. 2020; Mirzabaev et al. 2023).

We took an ecological perspective and started tracing mechanisms from productivity increases through the food system. One could equally begin the analysis with a socioeconomic view and start from market constraints, where demand structures mean that any productivity gains will necessarily be redirected toward expandable sectors. These approaches are not necessarily equivalent and may reveal different aspects of the same underlying mechanisms. An important point is that physiological, management and technology, and broader socioeconomics are all tied to each other, and depending on the scale, a simple perturbation of one element—in this case productivity—can lead to significant restructuring of food systems with implications for global food security.

Our ESM-IAM coupling approach has several limitations that present opportunities for future research. First, our unidirectional coupling lacks feedback from human systems to climate, though recent bidirectional frameworks like E3SM-GCAM (Di Vittorio et al. 2025) offer promising research directions. Second, we focus on single-model projections using one ESM (GFDL-ESM4) and one IAM rather than model ensembles, potentially introducing climate-specific biases. While ensemble studies would be very challenging due to various processes and parameterizations in both ESMs and IAMs (Krey et al. 2019; van Beek et al. 2020), they could provide broader uncertainty ranges and valuable insights, though caution is needed since IAMs can differ fundamentally in socioeconomic assumptions, meaning coupled approaches may not always allow direct comparisons. Third, while crop-specific models might seem more relevant and there are efforts to couple them with IAMs (Ruane et al. 2017; Filatova et al. 2025), our interest in vegetation productivity across all land types necessitated the broader CLASSIC framework.

Finally, we intentionally limited our analysis to productivity changes to maintain our counterfactual focus, though future work incorporating additional climate variables and their interactions would enhance understanding (Luo et al. 2025).

## 5 Conclusion

Our analysis reveals that climate-driven vegetation productivity increases do not simply translate to food abundance but instead fundamentally reorganize global food systems. By isolating the vegetation productivity signal from confounding management and technological factors, we demonstrate that productivity gains trigger systematic reorganization rather than simple expansion of food systems. The collapse of sugar crops and corn production alongside surges in soybeans and oil crops reflects systematic reallocation toward sectors with expandable demand, particularly bioenergy and animal feed. There is an extreme regional divergence such as Brazil’s massive production losses contrast sharply with gains in regions like EU-15 and Canada. These results highlight a critical gap in current food system projections: models that hold vegetation productivity constant or focus solely on crop yield responses miss the broader ecosystem competition that reshapes agricultural landscapes as climate change accelerates vegetation productivity across all terrestrial ecosystems, not just croplands. Understanding these reorganization patterns—rather than assuming simple productivity-to-food translations—is essential for anticipating how climate change will reshape not just how much food we produce, but what we produce, where we produce it, and ultimately who has access to it.

