## Supplementary Information for "Climate-Driven Ecosystem Productivity Changes Restructure Food Systems"

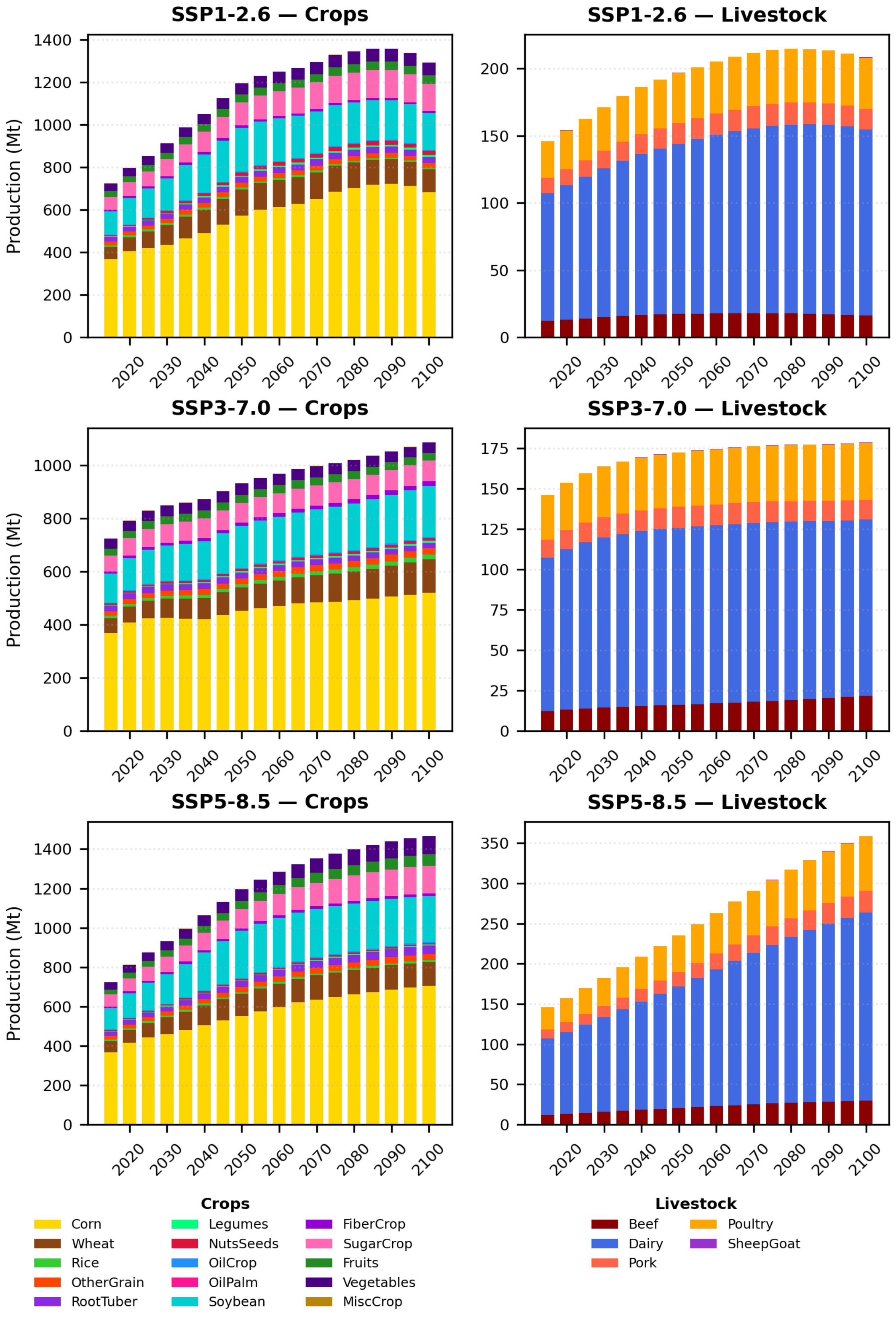


Figure S1. Global agricultural and livestock production, 2015-2100. Stacked barplot showing total production (Mt) for agricultural crops and livestock across scenarios.

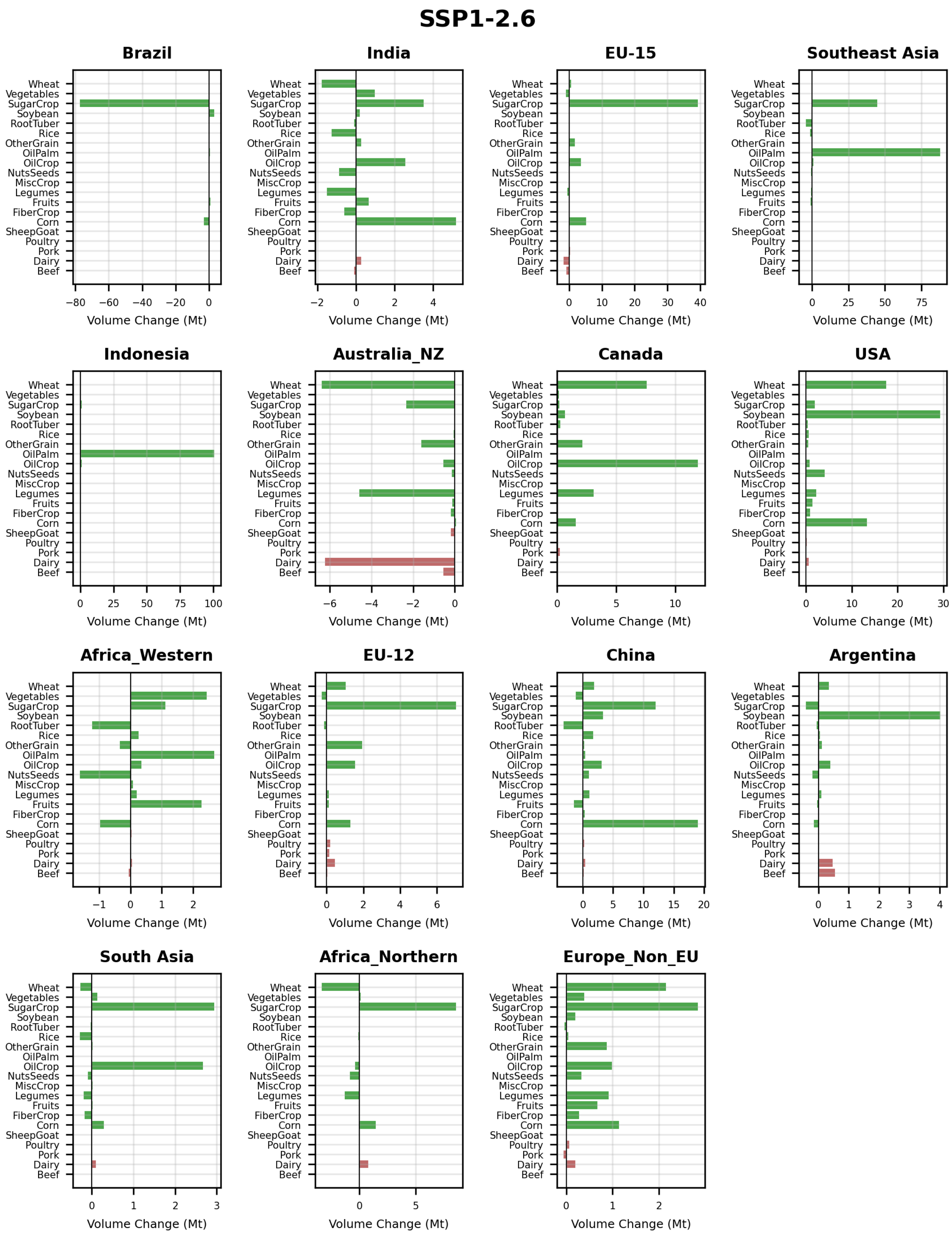


Figure S2. **Regional food item volume changes due to increased vegetation productivity.** Horizontal bar charts showing absolute changes (Mt) in individual food items for the 15 most responsive regions. Brown bars represent livestock products, green bars represent crop products. Results shown for each SSP scenario.

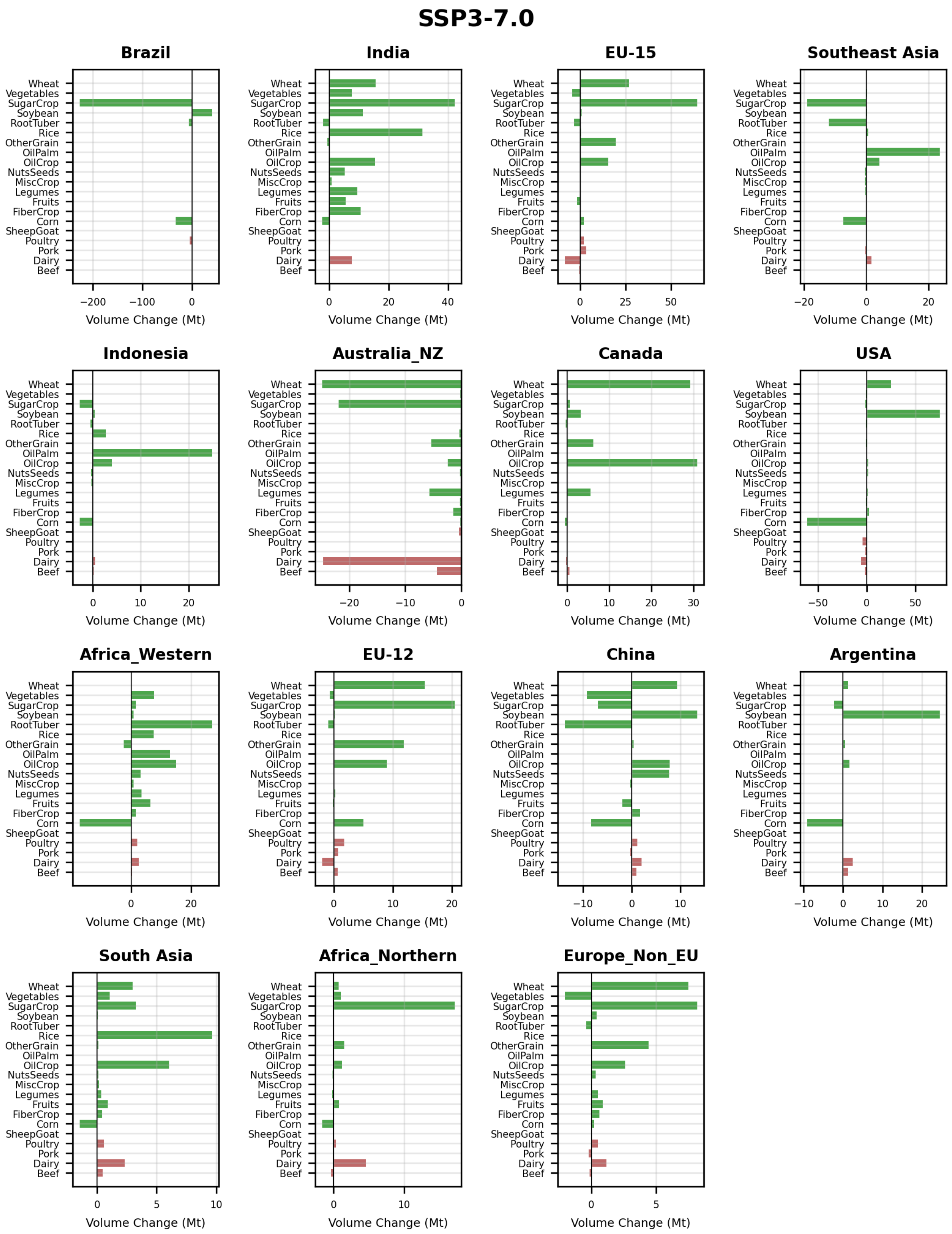


Figure 2 (continued).

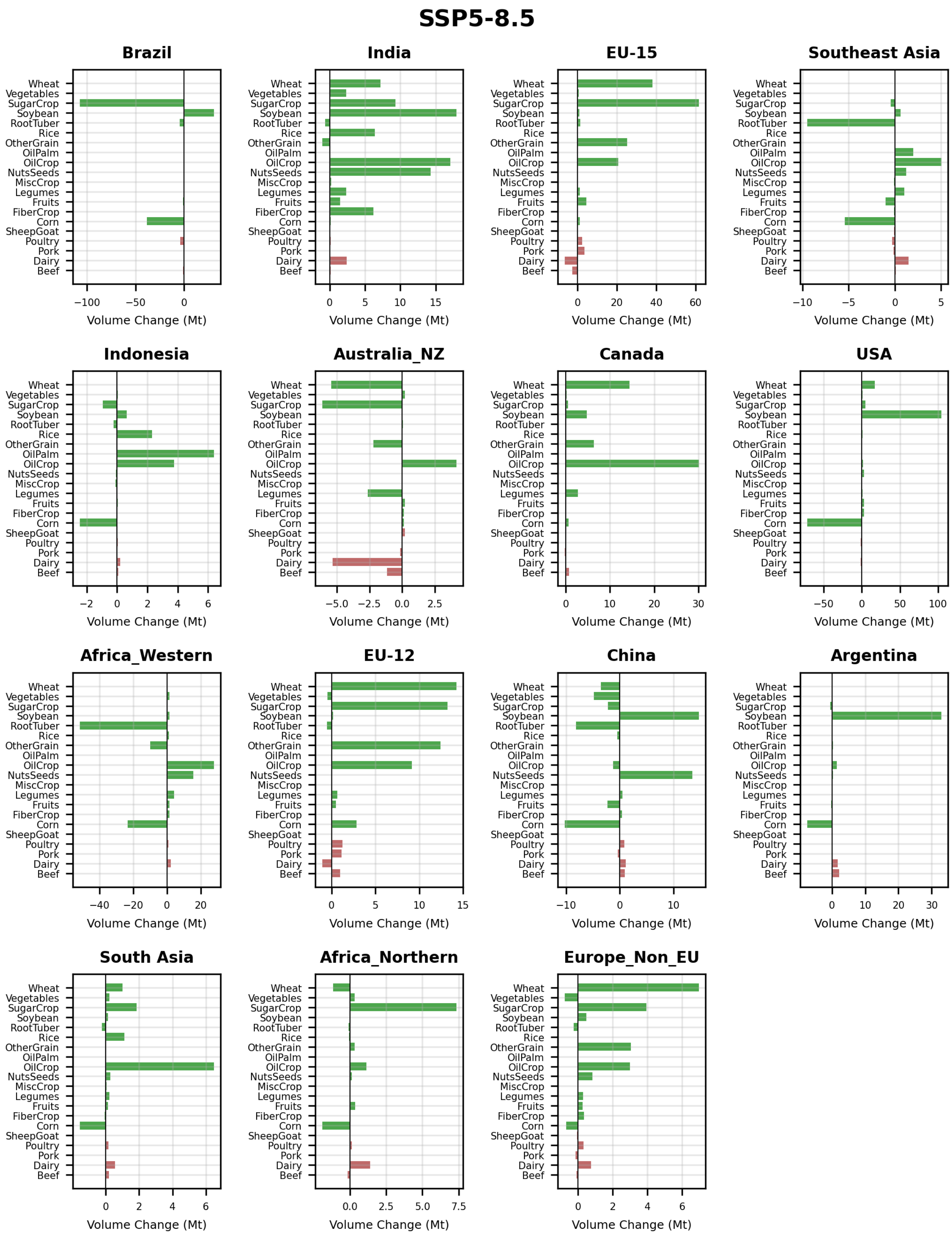


Figure 2 (continued).

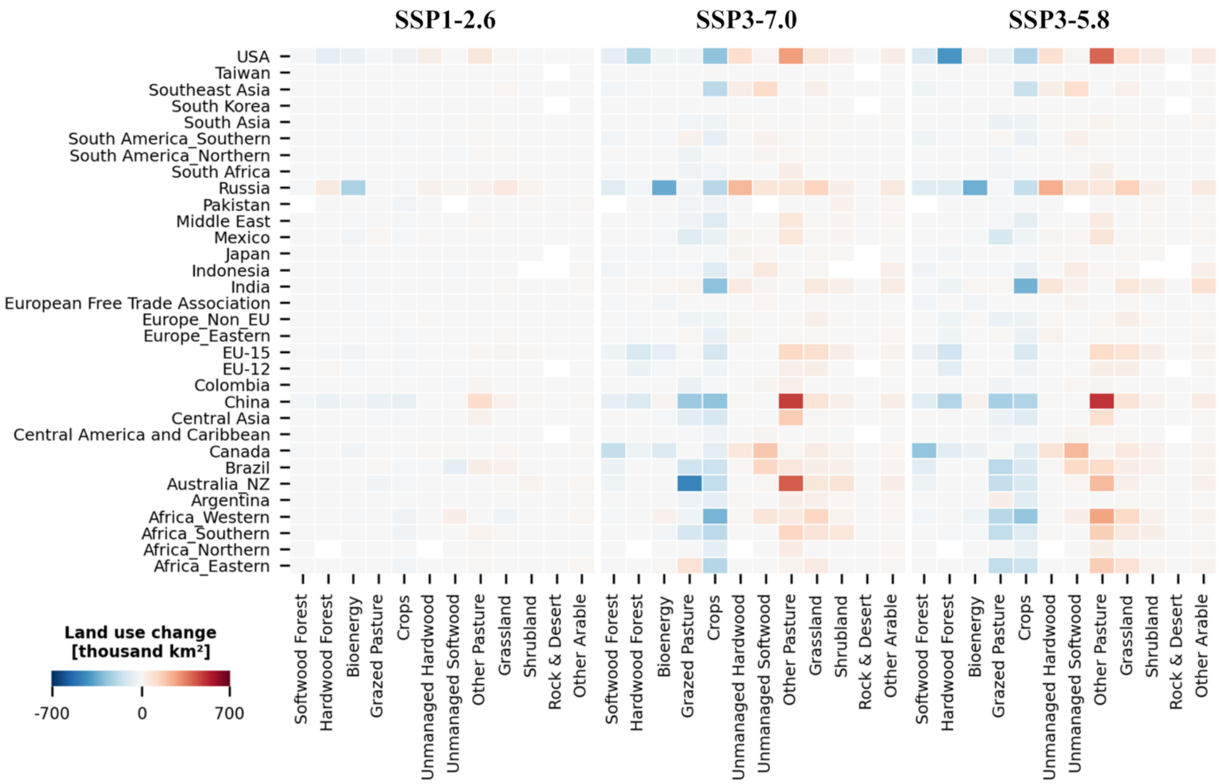


Figure S3. **Regional land use changes due to increased vegetation productivity.** Heatmaps showing land use change (thousand $km^2$) across regions and land use categories for SSP1-2.6, SSP3-7.0, and SSP5-8.5 scenarios.
